# Rapid Determination of Antibiotic Susceptibility by Stimulated Raman Scattering Imaging of D_2_O Metabolism

**DOI:** 10.1101/496778

**Authors:** Weili Hong, Ji-Xin Cheng

## Abstract

Rapid antibiotic susceptibility testing (AST) is urgently needed for treating infections with correct antibiotics and slowing down the emergence of antibiotic resistant bacteria. Current clinical methods reply on culture and take at least 16 h. Here, using *P. aeruginosa*, *E. coli and S. aureus* as models, we show that the AST can be finished in 10 minutes by stimulated Raman scattering (SRS) imaging of D_2_O metabolic activities. The metabolic incorporation of D_2_O, which is used for biomolecule synthesis, can be monitored in a single bacterium. Time lapse experiments show that the C-D vibrational signal can be observed in a single bacterium within 10 minutes culture in D_2_O medium. Since water is universally used for biosynthesis in bacteria, SRS imaging of D_2_O metabolism has the potential to be generalizable to different bacteria species.

---

Antimicrobial resistance has become a growing public threat, causing nearly 1 million deaths from drug-resistant infections each year globally^1^. It was estimated that by 2050, antimicrobial resistance will cause 10 million deaths and $100 trillion GDP loss if no action is taken^1,2^. To combat this crisis, new rapid diagnostic methods for antibiotic susceptibility testing (AST) are essential to reduce the deaths caused by drug-resistant infections^3^, and slow down the emergence of antimicrobial resistance. The culture-based phenotypic method remains the gold standard for AST. However, this method is too slow for guidance of immediate decision for infectious disease treatment. Genotypic methods, which detect known resistance genes, can provide faster results, but are not generally applicable to different bacteria and mechanism of resistance^4^.

To overcome these limitations, other methods have been developed for rapid AST, including, for example, microfluidic devices that increase the detection sensitivity by confining the sample in a small area^5–10^, imaging-based methods that monitor the growth or morphology change at the single cell level^10–12^, nucleic acids-based phenotypic AST quantifying nucleic acids copy number with antibiotic treatment^4,13,14^, and monitoring the spectral response to antibiotic treatment by Raman spectroscopy^15^. While these methods significantly reduce the time for AST, they either are not generalizable to different bacteria, or rely on cell proliferation, which has difficulties for viable but nonculturable species^16^, or require gene marker information before testing^17^.

We have previously demonstrated that the AST of bacteria can be accomplished within one cell cycle (30 minutes) by measuring the metabolic activity in single bacteria under a stimulated Raman scattering (SRS) microscope^18^. Specifically, we monitored the metabolic activity of deuterated glucose, glucose-d_7_, with chirped picosecond SRS at C-D vibrational frequency and used the C-D signal as a marker to perform AST. Like glucose, water is also ubiquitously used for biomolecule synthesis in bacteria^19^, and its metabolism can be selectively probed via monitoring the conversion of heavy water (D_2_O) into deuterated biomolecules at C-D vibrational frequency. Unlike glucose-d_7_, D_2_O itself does not have C-D bonds, therefore providing a better contrast for SRS metabolic imaging. The metabolic activity of D_2_O has been used to study the metabolic - active bacteria with spontaneous Raman spectroscopy^16,19^. However, the speed of spontaneous Raman is limited by the weak Raman scattering process^20^. Compared to spontaneous Raman, SRS has orders-of-magnitude signal enhancement, thereby enabling high speed chemical imaging of single cells ^21–26^. SRS imaging of D_2_O metabolism was recently demonstrated to be a noninvasive method to visualize metabolic dynamics in mammalian cells and live animals^27^.

Here, we demonstrate that femtosecond SRS imaging of D_2_O metabolism can determine the susceptibility of bacteria within 10 minutes. We show that the metabolic activity of D_2_O can be monitored in single *P. aeruginosa*, *E. coli* and *S. aureus*. For the broad C-D vibrational spectrum, the SRS signal of bacteria can be improved by more than 5 times by femtosecond SRS compared to the chirped SRS. The D_2_O metabolism in bacteria responds differently as fast as 10 minutes to different antibiotics, depending on the susceptibility of bacteria. Our work shows the promise of using SRS microscopy for metabolic activity studies and rapid AST at single bacterium level.

## Experimental Section

### SRS microscope

A femtosecond (fs) pulsed laser (InSight DeepSee, Spectra-Physics) with an 80-MHz repetition rate and dual outputs was employed for the SRS microscope. One 120 fs laser with tunable 680–1100 nm wavelength was served as the pump beam. The other 220 fs laser centered at 1040 nm, served as the Stokes beam, was modulated by an acousto-optical modulator (AOM, 1205-C, Isomet) at ~2.4 MHz. The two beams were collinearly combined through a dichroic mirror. When spectral focusing is needed for hyperspectral SRS, both beams were chirped with two 15 cm long SF57 glass rods. After chirping, the pulse durations of the pump and Stokes laser were 1.9 ps and 1.3 ps, respectively. For implementation of SRS imaging with femtosecond pulses, the glass rods were removed from the path. The pump and Stokes beams were directed into a lab-built laser scanning microscope. A 60x water objective (NA=1.2, UPlanApo/IR, Olympus) was used to focus the lasers to the sample, and an oil condenser (NA=1.4, U-AAC, Olympus) was used to collect the signal from the sample. Two filters (HQ825/150m, Chroma) were used to filter out the Stokes beam, the pump beam was detected by a photodiode (S3994-01, Hamamatsu) and the stimulated Raman signal was extracted by a lock-in amplifier (HF2LI, Zurich Instrument).

### Sample preparation

To make D_2_O containing LB medium, D_2_O was first mixed with purified water, then LB broth powder (Sigma Aldrich) was added to the solution with a final concentration of 2% in weight. The solution was sterilized by filtering. To prepare bacteria samples, *E. coli*, *S. aureus* or *P. aeruginosa* were cultivated in normal LB medium for 2-3 h to reach to log phase, then bacteria were diluted in a 1:100 ratio to the D_2_O containing LB medium. After incubation for a controlled period, 500 μl sample was centrifuged, washed twice with purified water, and deposited to an agarose gel pad.

To prepare the agarose gel pad, ~1% in weight agarose powder was added to 5 ml purified water in a plastic tube, then the tube was heated in microwave for about 20 seconds to melt the agarose powder. About 10 μl heated agarose gel solution was added to a coverglass by a pipette, another coverglass was immediately put on the top of the agarose gel solution to make it flat. After ~2 min, one coverglass was removed from the agarose gel by sliding the two coverglasses. Bacteria in solution were deposited to the gel pad, then another coverglass was put on top of the gel pad for SRS imaging.

### SRS imaging

To image bacteria at the C-D region, the pump wavelength was tuned to 849 nm, and the power at the sample was ~12 mW; the Stokes wavelength was fixed at 1040 nm, and the power at the sample was ~90 mW. Each image contains 200 × 200 pixels, the pixel dwell time is 30 μs.

### Spontaneous Raman spectroscopy

Bacteria in solution were deposited on a coverglass for spontaneous Raman measurement. Spontaneous Raman spectra of bacteria were acquired with an inverted Raman spectrometer (LabRAM HR evolution, Horiba scientific) with 532 nm laser source. The laser power at the sample was ~12 mW after 40x air objective, the acqusition time was 10 s. The grating was 600 l/mm.

### Broth dilution method

Bacteria were cultured in D_2_O containing LB medium in a 96-well plate. Antibiotics, using triplicate samples, were added to the plate and serially diluted. After about 20 h incubation at 37 °C, plates were visually inspected, and the MIC was categorized as the concentration at which no visible growth of bacteria was observed.

## Results and Discussion

### D_2_O induces negligible toxicity to bacteria

We first tested whether D_2_O has toxicity to bacteria by measuring their growth in D_2_O containing medium. Three types of bacteria (*E. coli*, *S. aureus*, and *P. aeruginosa*) were cultured at different concentrated D_2_O containing LB medium, and their growth were monitored with optical density (OD) measurement at 600 nm wavelength. We found that D_2_O concentration up to 100% did not show significant toxicity to the growth of *E. coli* and *S. aureus*, as indicated by the growth curve in D_2_O media of various concentrations (**Figure 1a** and **1b**). The growth of *P. aeruginosa* was initially slowed down in medium with D_2_O concentration of 70% and up, but eventually restored to normal growth after about 18 h for D_2_O concentration of 70% and 80%, and about 22 h for D_2_O concentration of 100% (**Figure 1c**). Therefore, D_2_O concentration of 70 % or lower in the medium does not induce significant toxicity to the bacteria.

**Figure 1.**
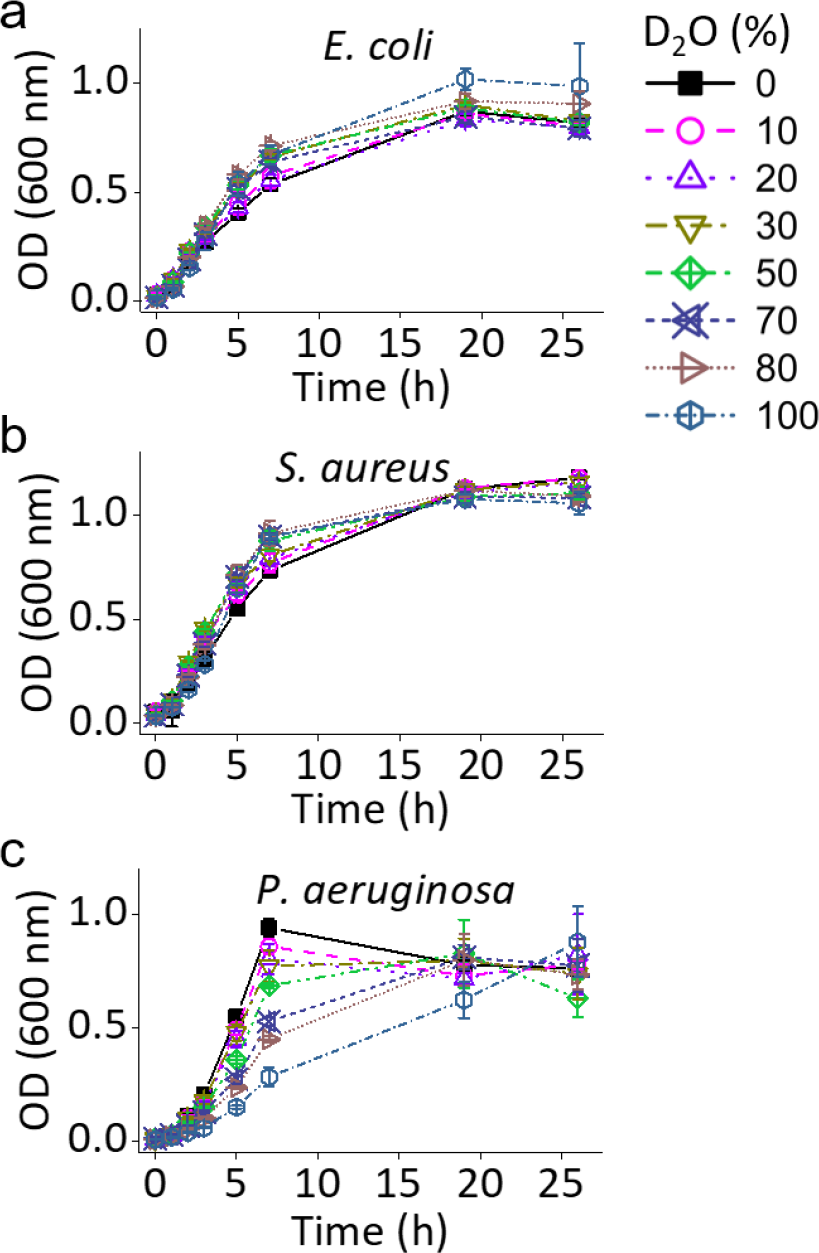
Testing of D_2_O toxicity to bacteria growth. Growth curve of *E. coli* (a), *S. aureus* (b) and *P. aeruginosa* (c) cultivated in LB medium with different D_2_O concentrations.

### Imaging metabolic incorporation of D_2_O in a single bacterium

We chose 70% D_2_O containing LB medium to cultivate bacteria, and used *P. aeruginosa* to test whether the D_2_O metabolic incorporation in a single bacterium can be monitored by our SRS microscope. Spontaneous Raman spectra of bacteria showed a broad peak (2070 2250 cm^−1^) at C-D vibrational region for bacteria cultivated in the D_2_O containing medium for 2 h (**Figure 2a**), indicating D_2_O had been successfully utilized for biomolecule synthesis. For control, bacteria cultivated in normal medium did not have this peak at this region (**Figure 2a**). To image single bacterium, bacteria were further diluted and deposited on an agarose gel pad. By tuning the Raman shift to C-D region (~2162 cm^−1^), a strong signal was observed for individual bacterium cultivated in D_2_O containing medium (**Figure 2b**, right). As a control, no C-D signal was observed for bacteria cultured in normal medium (**Figure 2b**, left). The results were confirmed by SRS spectra (**Figure 2c**) obtained through temporal tuning of chirped pump and Stokes femtosecond pulses. Similar results were obtained on *E. coli* (**Supporting Figure 1**) and *S. aureus* (**Supporting Figure 2**).

**Figure 2.**
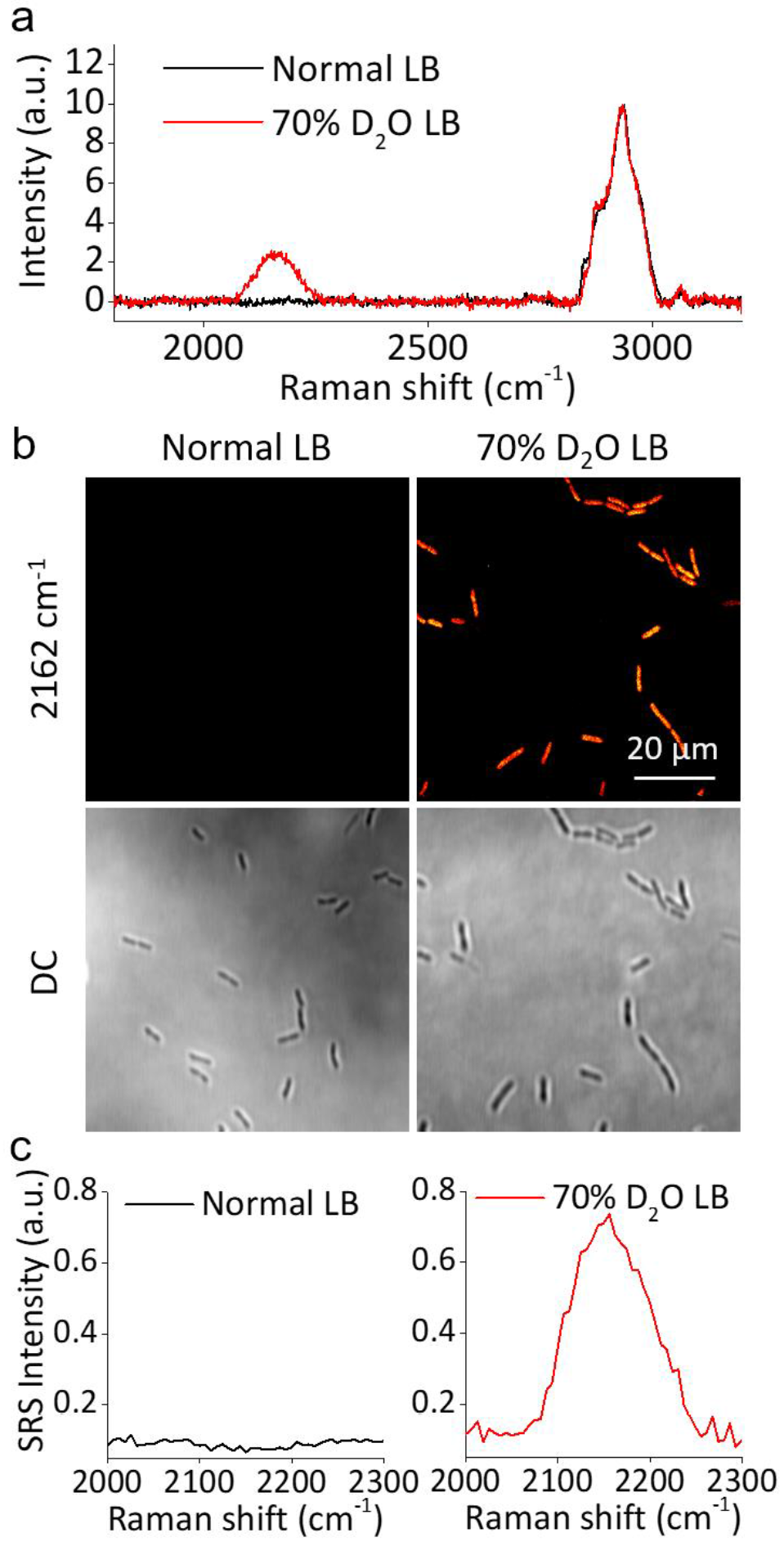
Biosynthesis in *P. aeruginosa* in D_2_O containing LB medium. a) Spontaneous Raman spectra of high-density *P. aeruginosa* cultivated in normal LB and 70% D_2_O containing LB medium for 2 h. b) SRS imaging at ~2162 cm^−1^ and the corresponding transmission images of *P. aeruginosa* cultivated in normal and 70% D_2_O containing LB medium for 2 h. c) Corresponding SRS spectra of a single bacterium in (b).

In order to shorten the D_2_O culture time, we tested whether non-chirped femtosecond pulses can enhance the signal over the chirped picosecond pulses. Because the C-D vibration band is relatively broad with a width of 180 cm^−1^ (**Figure 2c**), we hypothesized that femtosecond SRS without chirping could significantly increase the signal to noise ratio (SNR). To test this, we cultivated *P. aeruginosa* in 70% D_2_O containing LB medium for 30 minutes, and imaged them at ~2162 cm^−1^ by chirped picosecond pulses and non-chirped femtosecond pulses, respectively (**Figure 3a** and **3c**). The pump and Stokes powers were adjusted to make sure the same average pump and Stokes powers were used. The SNR of individual bacterium with picosecond and femtosecond SRS was 1.43 and 7.81, respectively, indicating ~5.5 times SNR improvement with femtosecond SRS over picosecond SRS (**Figure 3b** and **3d**). This improvement is attributed to two factors. First, the C-D vibrational band is broad, and the femtosecond SRS can detect broader band signal than the chirped picosecond SRS. Second, the pulse chirping reduced the peak power of the pulses. Although the same average power was used at the sample, the reduced peak power decreased the signal level due to the nonlinear effect of SRS.

**Figure 3.**
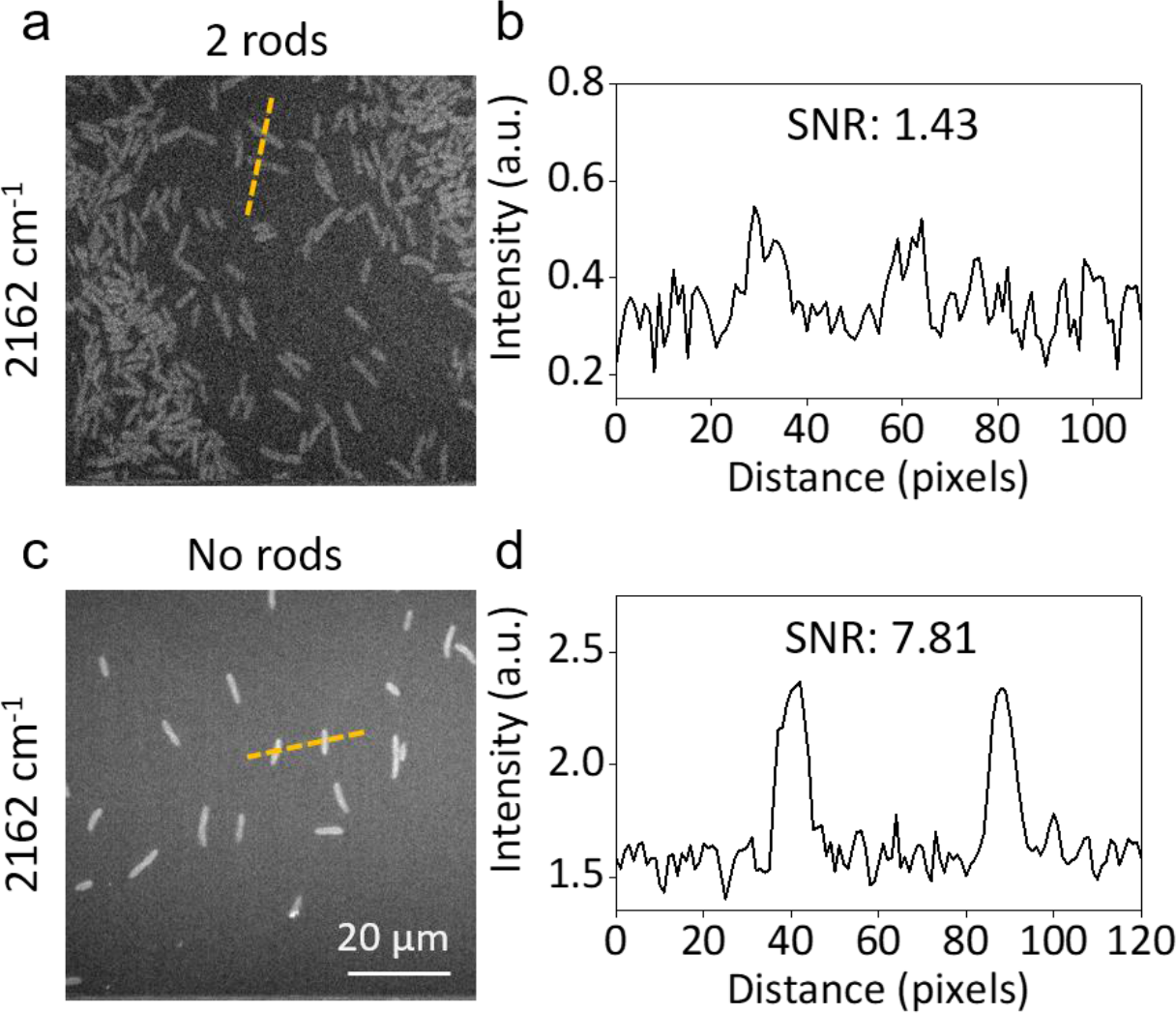
Femtosecond SRS improves SNR at C-D vibrational region over the chirped SRS. a) SRS imaging at ~2162 cm^−1^ of *P. aeruginosa* cultivated in 70% D_2_O containing LB medium for 30 minutes with picosecond pulses generated by chirping with two SF57 glass rods. b) Intensity plot of the orange line over bacteria in (a). c) SRS imaging at ~2162 cm^−1^ of *P. aeruginosa* cultivated in 70% D_2_O containing LB medium for 30 minutes with non-chirped femtosecond pulses. d) Intensity plot of the orange line over bacteria in (c).

### Time lapse measurement of D_2_O metabolic incorporation in a single bacterium

Next, we studied the time lapse of D_2_O metabolic activity in single *P. aeruginosa* with femtosecond SRS. The *P. aeruginosa* was cultivated in 70% D_2_O containing LB medium for up to 3 h. Figure 4a shows the SRS images of single *P. aeruginosa* at ~2162 cm^−1^, C-D signal in individual *P. aeruginosa* can be observed after culture as short as 10 minutes. Statistical analysis showed that the average C-D signal intensity in individual bacterium increases with time, and saturates at ~1.5 h (**Figure 4c**), which is about three generations since the generation time of *P. aeruginosa* cultivated in LB medium is 24-27 minutes^28^. To view individual bacterium more clearly, the images in **Figure 4a** were further zoomed in (**Figure 4b**). Interestingly, in the 10 minutes result, a stronger signal was observed in the cell periphery of bacterium (**Figure 4b**), as indicated by the intensity plot over the bacteria (**Figure 4d**). In contrast, in and after 30 minutes, the signal intensity is stronger in the intracellular area (**Figure 4b**), as indicated by the intensity plot over bacteria in the 30 minutes result (**Figure 4e**). Collectively, these results suggest that D_2_O is initially used to synthesize cell membrane and/or cell wall in *P. aeruginosa*.

**Figure 4.**
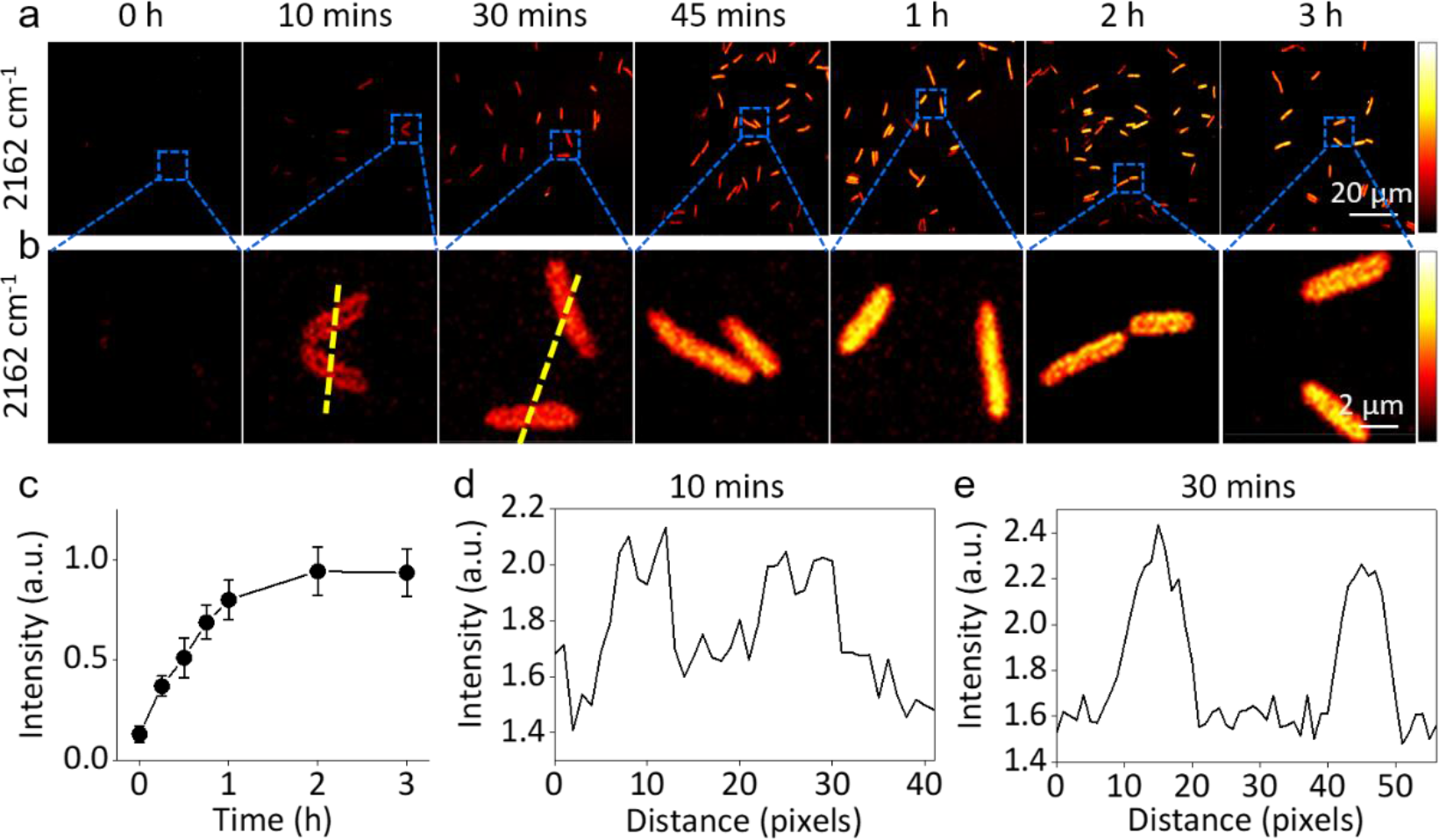
Time lapse SRS imaging of D_2_O-based biosynthesis in *P. aeruginosa*. a) SRS imaging of *P. aeruginosa* cultivated in 70% D_2_O containing LB medium from time 0 to 3 h. b) Zoom in SRS imaging of individual *P. aeruginosa* in the rectangular square in (a). c) Growth of average SRS intensity of individual *P. aeruginosa* shown in panel a. Error bars indicate standard deviation (N > 10). d) Intensity plot of the yellow line over bacteria from the 10 minutes result in (b). e) Intensity plot of the yellow line over bacteria from the 30 minutes result in (b).

### Rapid AST

To examine how antibiotics affect the metabolic incorporation of D_2_O in bacteria, and whether it can be used for rapid AST through SRS imaging, *P. aeruginosa* were cultivated in 70 % D_2_O containing LB medium, with the addition of 20 μg/ml gentamicin or cefotaxime. The susceptibility of *P. aeruginosa* were pre-determined to be susceptible to gentamicin and resistant to cefotaxime at this concentration by the conventional culture-based microdilution method. SRS imaging at ~2162 cm^−1^ showed that the C-D signal was significantly reduced after cultivation in 20 μg/ml gentamicin (**Figure 5a**), indicating that the metabolic activity of D_2_O in *P. aeruginosa* was inhibited by gentamicin. On contrary, *P. aeruginosa* cultivated in cefotaxime can be observed at ~2162 cm^−1^ SRS imaging at all time points, indicating active metabolic incorporation of D_2_O in *P. aeruginosa* when cultivated in cefotaxime. We observed that *P. aeruginosa* tends to form long rods when cultivated in cefotaxime (**Figure 5b**). This filamentary formation, which happens when Gram-negative bacteria are treated with β-lactam antibiotics, was also observed for *P. aeruginosa* treated with ceftazidime^10^.

**Figure 5.**
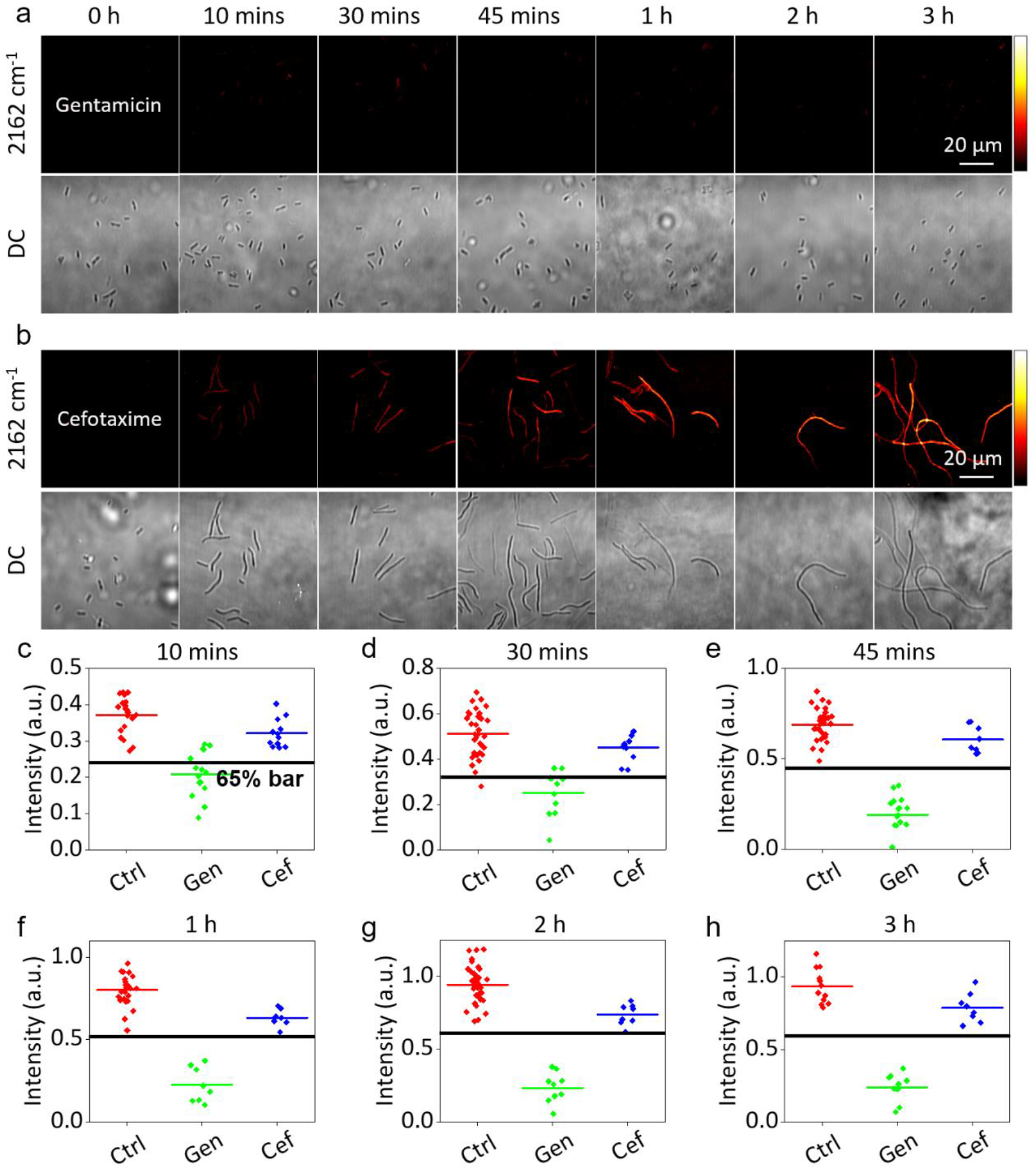
SRS-based antibiotic susceptibility testing of *P. aeruginosa*. a, b) C-D SRS imaging at ~2162 cm^−1^ and transmission images of *P. aeruginosa* cultivated in 70% D_2_O containing LB medium with the addition of 20 μg/ml gentamicin (a) or cefotaxime (b). c - f) Average C-D signal intensity of *P aeruginosa* cultivated in 70% D_2_O containing LB medium without antibiotic treatment (control), and with gentamicin or cefotaxime treatment for 10 minutes (c), 30 minutes (d), 45 minutes (e), 1 h (f), 2 h (g) or 3 h (h). The colour bar indicates average C-D signal intensity of individual *P. aeruginosa* in each group. Black bar indicates the 65% intensity value of the average *P. aeruginosa* C-D signal intensity in control.

To examine whether the D_2_O metabolic activity of bacteria can be used to rapidly differentiate the antibiotic susceptibility of bacteria, the average C-D signal intensity of bacteria was compared between three groups (**Figure 5c - 5h**), the control without antibiotics treatment (**Figure 4a**), treated with gentamicin (**Figure 5a**), and treated with cefotaxime (**Figure 5b**). To separate the susceptible and resistant group, we determined a 65% line threshold, which is 65% the average C-D intensity of bacteria in control, in all plots from 10 minutes to 3 h results (**Figure 5c - 5h**). This threshold can clearly divide the susceptible and resistant groups, the group treated with gentamicin was always below the threshold, and the group treated with cefotaxime was always above the threshold. Therefore, the susceptibility of *P. aeruginosa* to gentamicin and cefotaxime can be determined in as short as 10 minutes.

### Metabolic activity based MIC determination

To test whether SRS metabolic imaging can quantitate the minimal inhibitory concentration (MIC) of antibiotics to bacteria, we cultivated *P. aeruginosa* in 70% D_2_O containing LB medium for 1 hour with the addition of gentamicin with serial diluted concentration. SRS imaging at ~2162 cm^−1^ showed that the D_2_O metabolic activity of *P. aeruginosa* was inhibited at 8 μg/ml or higher concentrated gentamicin cultivation (**Figure 6a**). For control, no C-D signal was observed for *P. aeruginosa* cultivated in normal LB medium. The average intensity of *P. aeruginosa* C-D signal was plotted and compared. With the 65% intensity threshold, the metabolic activity based MIC was determined to be 8 μg/ml by the SRS-based D_2_O metabolic imaging method (**Figure 6b**). This value is consistent with the MIC determined by the broth dilution method.

**Figure 6.**
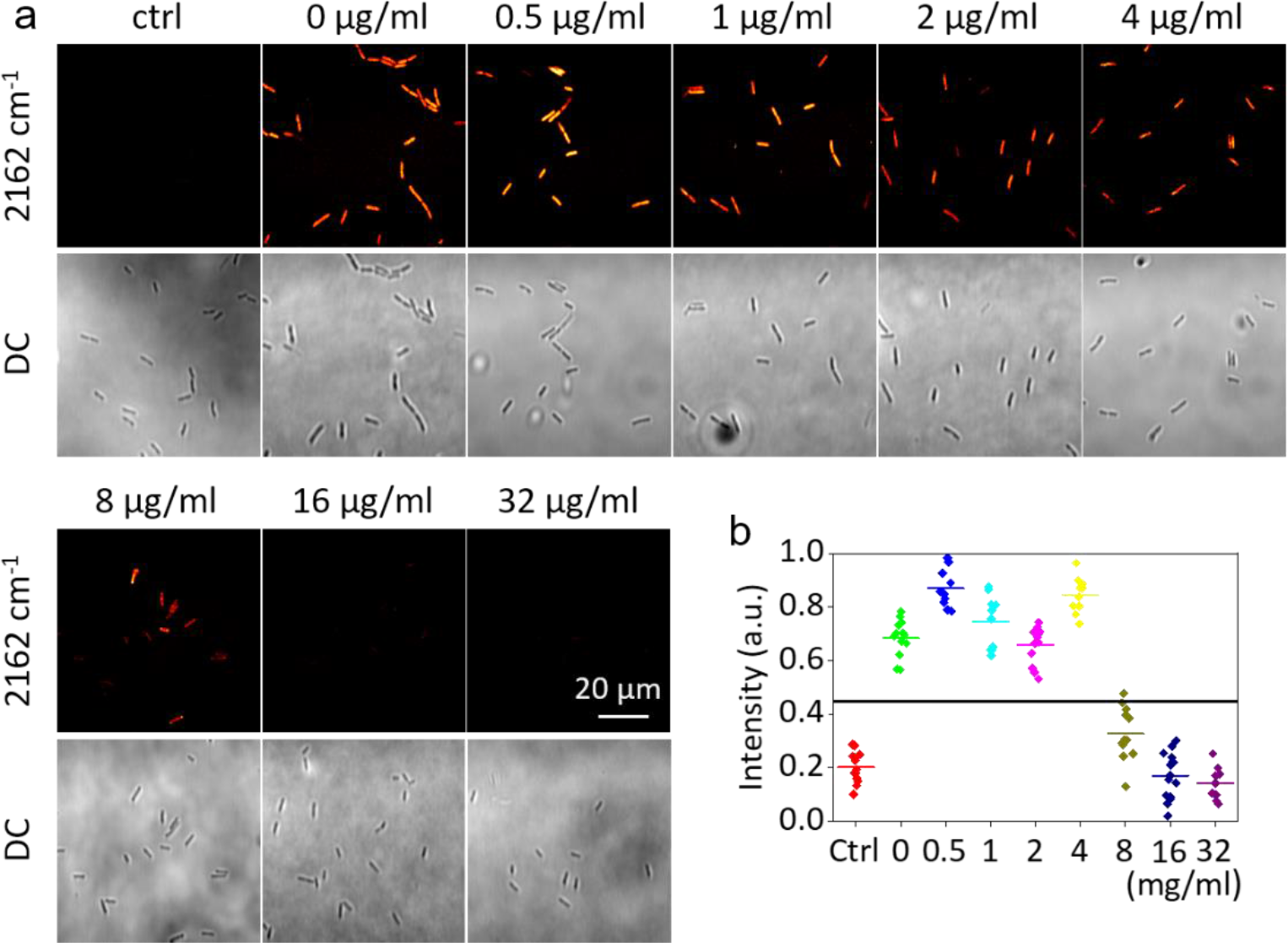
MIC determination of *P. aeruginosa* against gentamicin. a) C-D SRS imaging at ~2162 cm^−1^ and transmission images of *P. aeruginosa* cultivated in normal LB (control) or 70% D_2_O containing LB medium with the addition of different concentrated gentamicin. b) Average C-D signal intensity of individual *P. aeruginosa* in (a). Black bar indicates 65% intensity bar of the average SRS intensity of individual *P. aeruginosa* in D_2_O containing LB medium without treatment (0 μg/ml).

## Conclusion

We demonstrated rapid determination of the susceptibility of bacteria in 10 minutes by SRS imaging of the D_2_O metabolic incorporation in bacteria. Femtosecond SRS imaging can monitor D_2_O metabolism in bacteria at the single cell level with high signal to noise ratio. Antibiotics can inhibit the metabolic activity of D_2_O when the bacteria are susceptible to this antibiotic, and this inhibition can be observed after culture in D_2_O containing medium for 10 minutes. The metabolic activity based MIC can be quantitated by our method. Because water is an essential molecule for biosynthesis in bacteria, the SRS-based D_2_O metabolic imaging method has the potential to be generalized for rapid AST in various species including clinical samples.

## AUTHOR INFORMATION

### Author Contributions

J.X.C, and W.H. conceived the idea. W.H. designed the experiment. W.H. and L.L. conducted the experiment. W.H. analyzed the data. W.H., and J.X.C. co-wrote the manuscript. All authors have given approval to the final version of the manuscript.

### Notes

The authors declare no competing financial interests.

## Supporting information

## ACKNOWLEDGMENT

This work was supported by Keck Foundation Science & Engineering Grant and R01GM118471 to JXC.

