## Supplementary material for "Rapid Determination of Antibiotic Susceptibility by Stimulated Raman Scattering Imaging of D_2_O Metabolism"

**Abstract:** Rapid antibiotic susceptibility testing (AST) is urgently needed for treating infections with correct antibiotics and slowing down the emergence of antibiotic resistant bacteria. Current clinical methods reply on culture and take at least 16 h. Here, using *P. aeruginosa*, *E. coli and S. aureus* as models, we show that the AST can be finished in 10 minutes by stimulated Raman scattering (SRS) imaging of $D_2O$ metabolic activities. The metabolic incorporation of $D_2O$, which is used for biomolecule synthesis, can be monitored in a single bacterium. Time lapse experiments show that the C-D vibrational signal can be observed in a single bacterium within 10 minutes culture in $D_2O$ medium. Since water is universally used for biosynthesis in bacteria, SRS imaging of $D_2O$ metabolism has the potential to be generalizable to different bacteria species.

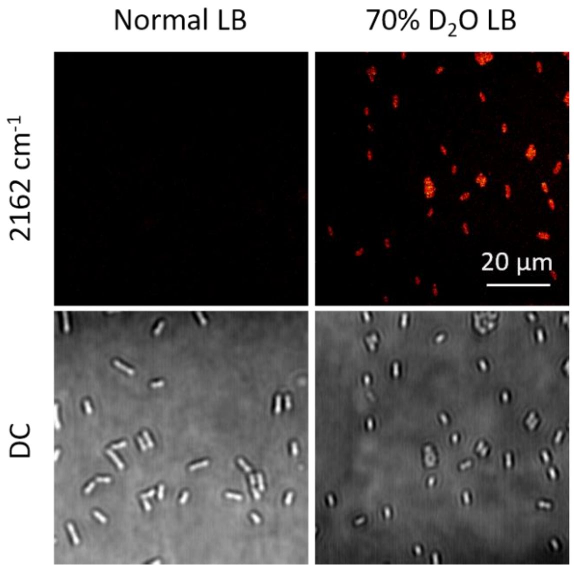


**Figure S1.** SRS imaging at ~2162 $cm^{-1}$ and the corresponding transmission images of *E. coli* cultivated in normal and 70% $D_2O$ containing LB medium for 2 h.

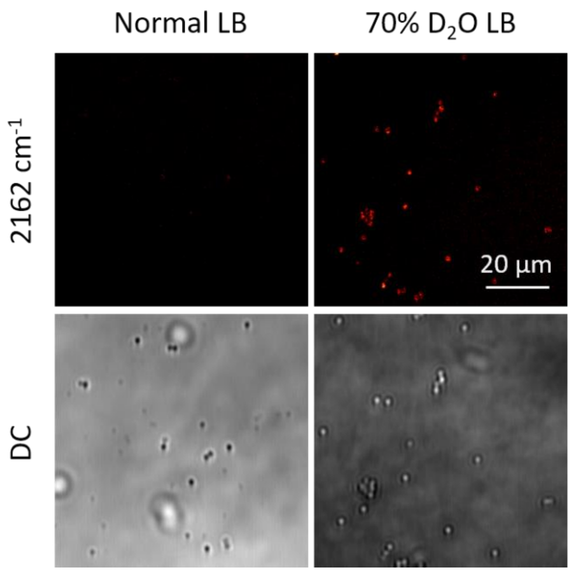


**Figure S2.** SRS imaging at ~2162 $cm^{-1}$ and the corresponding transmission images of *S. aureus* cultivated in normal and 70% $D_2O$ containing LB medium for 1 h.
